# Integrin-Binding Matricellular Protein Fibulin-5 Maintains Epidermal Stem Cell Heterogeneity During Skin Aging

**DOI:** 10.1101/2025.05.26.656217

**Authors:** Wenxin Fan, Mizuho Ishikawa, Erna Raja, Ahmed M. Hegazy, Hinata Date, Yen Xuan Ngo, Yoshifumi Sato, Kazuya Yamagata, Hiromi Yanagisawa, Aiko Sada

**Author notes:** Correspondence addressed to: Aiko Sada, Ph.D. Division of Skin Regeneration and Aging, Medical Institute of Bioregulation, Kyushu University, 3-1-1 Maidashi, Higashi-ku, Fukuoka City, Fukuoka, 812-8582, Japan.

## Abstract

The extracellular matrix (ECM) is crucial in building the extracellular environment and translating extracellular information into biochemical signals that sustain organ functions. Fibulin-5 (*Fbln5*) is a multifunctional ECM protein essential for forming elastic fibers and regulating cellular functions by binding to integrins. While fibulin-5 expression decreases with age in human skin, the functional implications of this decrease, particularly in epidermal stem cell regulation, have remained largely unexplored. Here, we find that the knocking out of *Fbln5* in mice results in early impairments of epidermal stem cell properties, similar to the chronological aging of the skin. The *Fbln5* deficiency suppresses the expression of integrins and other cell junction proteins, leading to the inactivation of YAP signaling in epidermal stem cells. This reduction in YAP signaling results in the downregulation of the fast-cycling epidermal stem cell marker SLC1A3 in human skin and primary keratinocytes. These results suggest that, in addition to its known function in elastic fiber formation, fibulin-5 also plays a critical role in regulating the balance of epidermal stem cell populations during skin aging and coordinating the crosstalk between the extracellular environment and intracellular signaling.

## 1. Introduction

The interfollicular epidermis, the outermost layer of the skin, functions as a protective barrier and is maintained by the proliferation and differentiation of epidermal stem cells throughout life. Recent single-cell and lineage analyses suggest that these stem cell populations are heterogeneous in their molecular properties and cellular lineages (Ghuwalewala et al., 2022; Joost et al., 2016; Koren et al., 2022; Mascré et al., 2012; Rognoni & Watt, 2018; Wang et al., 2020). Notably, mouse tail skin exhibits orthokeratotic interscale and parakeratotic scale structures, maintained by slow- and fast-cycling epidermal stem cell populations characterized by Dlx1 and Slc1a3, respectively (Gomez et al., 2013; Sada et al., 2016). During skin aging, there is a gradual loss of Slc1a3^+^ fast-cycling epidermal stem cells, resulting in an imbalance of the epidermal stem cell population (Raja et al., 2022). Similar epidermal stem cell heterogeneity has been observed in human skin, with their localization corresponding to the undulating structures known as inter-ridges (dermal papillae) and rete ridges (Ghuwalewala et al., 2022; Lawlor & Kaur, 2015). However, the factors regulating epidermal stem cell heterogeneity in aging skin remain poorly understood.

The extracellular matrix (ECM) provides both a biochemical and mechanical environment for tissue stem cells. In the skin, the ECM includes the dermal compartment, primarily composed of collagen types I and III and elastic fibers, and the basement membrane zone at the epidermal-dermal junction (Chermnykh et al., 2018; Raja et al., 2023). Integrins, hemidesmosomes, and other cell junctional molecules anchor epidermal stem cells to the basement membrane, maintaining their undifferentiated state (Liu et al., 2019; Wang et al., 2022; Watt, 2002). Integrins also mediate the conversion of extracellular signals into intracellular signals, controlling the cellular function of epidermal stem cells (AlDahlawi et al., 2006; Morgner et al., 2015; Watt, 2002). Deleting integrin β1 in mice led to a loss of the parakeratotic scale area (López-Rovira et al., 2005), and aged human skin showed reduced integrin β1 expression in epidermal stem cells (Giangreco et al., 2010). Additionally, levels of fibulin-7 (*Fbln7*), a matricellular protein at the basement membrane, decrease in aged skin, and *Fbln7* knockout (KO) results in the loss of fast-cycling epidermal stem cells (Raja et al., 2022). Changes in ECM composition and structure alter the mechanical environment for hair follicle and epidermal stem cells, thereby modulating tissue and cellular functions (Aragona et al., 2020; Ichijo et al., 2022; Koester et al., 2021; Li et al., 2022; Xie et al., 2022). However, the factors coordinating the extracellular environment and intracellular signals to regulate epidermal stem cell heterogeneity remain elusive.

Fibulin-5 (*Fbln5*) is a structural component of an elastic fiber network (Yanagisawa et al., 2009). In the skin, fibulin-5 is localized in the papillary dermis, oriented perpendicular to the epidermis, and integrates with other elastic fiber components in the reticular dermis (Kadoya et al., 2005). During aging, fibulin-5 expression decreases, particularly in the papillary dermis region (Kadoya et al., 2005; Langton et al., 2019). *Fbln5* KO mice showed defects in elastic fiber formation, resulting in skin, arterial, and lung abnormalities (Yanagisawa et al., 2002). In individuals with cutis laxa, *FBLN5* is associated with the development of loose and sagging skin, resembling the appearance of premature aging (Gharesouran et al., 2021). Fibulin-5 has a role in the regulation of tissue stiffness and the inflammatory response in skin fibrosis (Nakasaki et al., 2015).

Fibulin-5 also facilitates cell surface integrin binding via its arginine-glycine-aspartic acid (RGD) motif (Budatha et al., 2011). Fibulin-5 directly interacts with integrins, including α5β1, αvβ3, and αvβ5 (Lomas et al., 2007; Nakamura et al., 2002; Yanagisawa et al., 2009). These integrins have been implicated in mechanotransduction, with α5β1 mediating adhesion strength and αvβ3 involved in the early stage of mechanotransduction (Roca-Cusachs et al., 2009). Given its interaction with these integrins, we hypothesize that fibulin-5 may influence epidermal stem cell regulation; however, this role remains underexplored. Therefore, this study investigates how the age-dependent decrease in fibulin-5 modulates the extracellular environment, affecting the cellular and molecular properties of epidermal stem cells. Our results suggest that fibulin-5 contributes to maintaining epidermal stem cell heterogeneity in aging skin, partly mediated through the integrin network and YAP-dependent intracellular signaling.

## 2. Materials and Methods

### 2.1. Mice and Ethics

Experimental mice were housed in specific pathogen-free facilities in the Center for Animal Resources and Development at Kumamoto University, the Laboratory of Embryonic and Genetic Engineering at Kyushu University, and the Laboratory Animal Resource Center at the University of Tsukuba. The mice were maintained under a 12/12-h light/dark cycle at a controlled temperature of 22°C ± 1–2 °C, with free access to water and standard chow. C57BL/6J mice were obtained from Charles River Laboratories Japan, Inc. (Kanagawa, Japan) and Japan SLC, Inc. (Shizuoka, Japan).

All animal procedures were conducted according to the institutional animal care guidelines of each university and were approved by the respective animal research and ethics committees. The generation of *Fbln5* KO mice (129SvEv; C57BL/6) was reported previously (Okuyama et al., 2017). BrdU (5-bromo-20-deoxyuridine; Sigma-Aldrich) was administered in drinking water (0.8 mg/mL) two days before sacrifice to label cells in the S-phase of the cell cycle. Mice of both sexes were used in the experiments and were randomly allocated to the groups.

### 2.2. Human Skin Samples and Ethics

Frozen full-thickness normal human abdominal skin samples representing the ages of 20–40 years (SK534, SK545, SK548, and SK550) and 50–60 years (PK003, SK0033, SK220, SK255, SK602, and SK0342) were purchased from CTI-Biotech (Lyon, France) under ethical considerations. Informed consent was obtained from anonymous donors to collect these samples, and the consent and collection procedures were compliant with European standards and applicable local ethical guidelines.

### 2.3. Whole Mount Immunostaining

Whole mount staining of mouse tail skin was performed as previously described (Sada et al., 2016). The following primary antibodies were used: rat anti-BrdU (1:300, Abcam, ab6326), mouse anti-K10 (1:100, Abcam, ab9026), guinea pig anti-K36 (1:100, Proteintech, 14309-1-AP), and rat anti-Ki-67 (1:100; eBioscience [Invitrogen], 14-5698-82). A mouse-on-mouse kit (Vector Laboratories) was used to detect the primary mouse antibody. Before incubation with the anti-BrdU antibody, the samples were treated with 2 N HCl at 37°C for 30 min. All samples were stained with Hoechst nuclear stain (Sigma-Aldrich) before mounting. The stained whole-mount epidermis was observed under a confocal microscope (Nikon A1 HD25 or Zeiss LSM900), and images were captured and analyzed using NIS Elements Imaging or Zen 3 (Blue Edition) software. The images were adjusted using Adobe Photoshop (version 2024). All whole-mount images were viewed as a projected Z-stack, viewed from the basal layer side.

### 2.4. Hematoxylin and Eosin Staining and Section Immunostaining

Mouse tail skin was directly embedded in an optimal cutting temperature compound (Tissue-Tek, Sakura). The 10 μm sections were fixed in 4% paraformaldehyde at room temperature (RT) for 10 min. Next, the sections were stained with hematoxylin (Wako) for 20 min and eosin Y (Wako) for 15 sec before dehydration and mounting in Entellan solution (Merck Millipore). Images were captured using an EVOS M5000 Imaging System (Thermo Fisher Scientific).

Skin sections were immunostained as previously described (Sada et al., 2016). The rabbit anti-fibulin-5 antibody (BSYN1923) was reported previously (Yanagisawa et al., 2002). The following primary antibodies were used: rabbit anti-collagen XVII (1:300; Abcam, ab184996), rat anti-integrin α6 (1:150; BD Bioscience, 555734), rabbit anti-integrin β1 (1:200; Cell Signaling Technology, 34971S), rabbit anti-integrin β3 (1:100; Thermo Fisher Scientific, MA5-32077), chicken anti-K5 (1:500; BioLegend, 905904), rat anti-Nectin-3 (1:100; Abcam, ab16913), guinea pig anti-K36 (1:100, Proteintech, 14309-1-AP), rabbit anti-YAP (1:100; Cell Signaling Technology, 4912S), rabbit anti-ASS1 (1:1000; Cell Signaling Technology, 05-665), and guinea pig anti-SLC1A3 (1:100; Frontier Institute, GLAST-GP-Af1000). The sections were counterstained with Hoechst (Sigma-Aldrich) for 10 min. The sections were imaged using a confocal microscope (Nikon A1 HD25 or Zeiss LSM900), and the images were adjusted using Adobe Photoshop (version 2024).

### 2.5. FACS Isolation

The subcutaneous and fat tissues were removed from the tail skin and incubated in 0.25% trypsin/EDTA overnight at 4°C and then at 37°C for 30 min the next day. Single-cell suspensions were prepared by gentle scraping of the epidermis. Next, the cells were stained with the following antibodies for 30 min on ice: biotin-conjugated CD34 antigen (1:50, eBioscience, 13-0341), APC-conjugated streptavidin (1:100, BD Biosciences, 554067), BUV395-conjugated integrin α6 (1:100, BD Biosciences, custom), and BV421-conjugated Sca-1 (1:100, BD Biosciences, 562729). Dead cells were excluded by 7-AAD staining (BD Biosciences). Cells were isolated using a FACS Aria flow cytometer (BD Biosciences). The data were analyzed using the FlowJo software (BD Biosciences).

### 2.6. RNA-Seq

The FACS-isolated cells from *Fbln5* WT and KO mice were directly sorted into Trizol (Ambion, 10296028) and submitted to Tsukuba i-Laboratory LLP at the University of Tsukuba. RNA integrity was analyzed using an Agilent 2100 bioanalyzer. The RNA-seq library was prepared using the SMARTer^®^ Stranded Total RNA-Seq Kit v2–Pico Input Mammalian (Takara) and sequenced using an Illumina NextSeq500 system. The data were analyzed using CLC Genomics 11 software (Qiagen). After normalization, genes exhibiting ≥2-fold differences were selected. Principal component analysis was performed using CLC Genomics 20.0 software. A volcano plot was generated using SRplot (Tang et al., 2023). Gene Ontology analysis was performed using Metascape (https://metascape.org). Hierarchical clustering was performed using Clustvis (https://biit.cs.ut.ee/clustvis/).

### 2.7. Cell Culture

Neonatal primary human keratinocytes (KER112002, KER112005, and KER112006; Biopredic International) were cryopreserved in CnT-Cryo-50 (CELLnTEC), and cells with fewer than five passages were used in the experiments. After thawing, keratinocytes were cultured in 100 mm dishes (7 mL medium/dish) pre-coated with collagen type IV (Sigma-Aldrich) in phosphate-buffered saline (PBS) and maintained in CnT-Prime medium (CELLnTEC) at 37°C in a humidified atmosphere with 5% CO^2^. Collagen type IV in PBS (1 mg/mL) was prepared at least 30 min before the experiments at RT. Once the cells reached 80%–90% confluency, they were passaged using Accumax (Innovative Cell Technologies) and Accutase (Innovative Cell Technologies).

For the cell density experiments, primary human keratinocytes were seeded at 150,000, 50,000, and 25,000 cells/well in 12-well plates. The plates were pre-coated with 50 μg/mL collagen type IV in PBS (Sigma-Aldrich) and maintained in CnT-Prime medium (CELLnTEC) for 48 h post-seeding under standard culture conditions. Samples were fixed with 4% paraformaldehyde (PFA) at RT for 10 min before staining. The keratinocytes were immunofluorescently stained as previously described (Xuan Ngo et al., 2021) using the following primary antibodies: rabbit anti-YAP (1:100; Cell Signaling Technology, 4912S), mouse anti-Ki-67 (1:200; BD Bioscience, 14-5698-82), rabbit anti-ASS1 (1:1000; Cell Signaling Technology, 05-665), and guinea pig anti-SLC1A3 (1:100; Frontier Institute, GLAST-GP-Af1000). The experiments were performed in biological triplicates unless otherwise stated in the figure legend.

### 2.8. RT-qPCR

Total RNA was isolated using an RNeasy Mini Kit (Qiagen). cDNA was synthesized using an iScript cDNA Synthesis Kit (Bio-Rad). RT-qPCR was conducted on a LightCycler 96 System (Roche) using the FastStart Essential DNA Green Master (Roche). A complete list of the primers used is provided in Supplementary Table 1.

### 2.9. Quantification and Statistical Analysis

All quantifications were performed independently in at least three independent experiments. The data are presented as the mean ± standard deviation (SD). Statistical analyses and graphs were generated using GraphPad Prism 9 software. *P*-values < 0.05 were considered statistically significant. The sample size was dictated by experimental considerations and not by a statistical method. The experiments were not randomized. The investigators were not blinded to allocation and outcome assessments during the experiments.

In tail sections, an epidermal unit was defined as the interfollicular epidermis region comprising one scale and one interscale structure between two hair follicles (Changarathil et al., 2019). The epidermal thickness was measured for at least six epidermal units per mouse using the ImageJ software (US National Institutes of Health; Fig. 1G, I; Fig. S1C). BrdU^+^ and Ki-67^+^ cells were manually counted from 4 – 8 projected Z-stack images per mouse, each containing 15 – 20 interscale or scale structures (Fig. 1L, M; Fig. S1E, G). K10^+^ and K36^+^ areas were measured in 6–8 projected Z-stack images per mouse, each containing 30 interscale or scale structures using the ImageJ software (Fig. 1O, Q, S; Fig. S1I; Fig. S2B). The integrated intensity of collagen XVII, integrin β1, integrin α6, and integrin β3 was measured in epidermal basal cells, excluding nuclear signals (Fig. 3D, F, H, J, L, N, P, R; Fig. S3C). At least 50 cells were analyzed for each sample.

**FIGURE 1.**
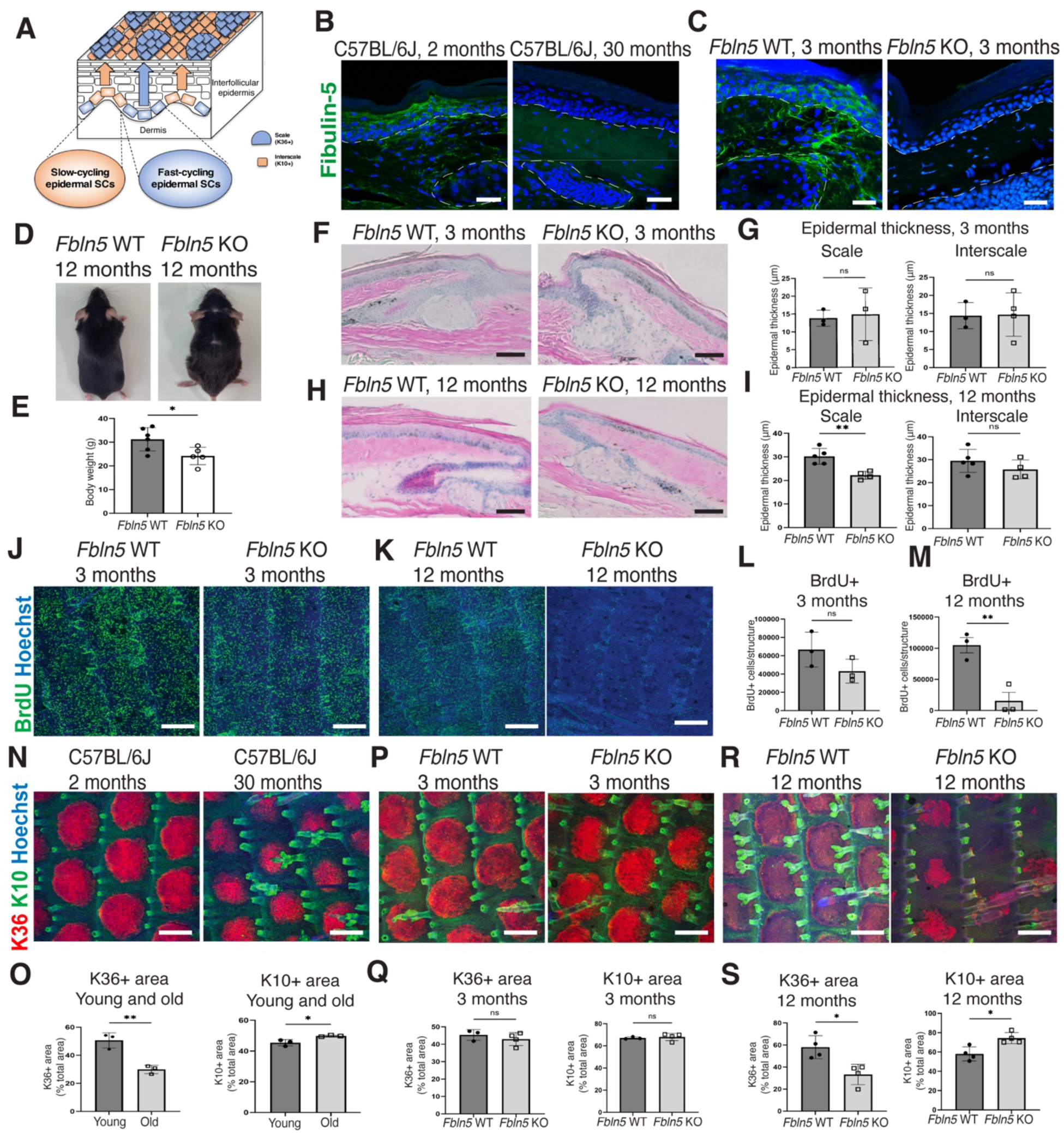
Impact of fibulin-5 deficiency on the skin aging process. (**A**) Schematic representation of the interfollicular epidermis of mouse tail skin. Slow-cycling epidermal stem cells (SCs) produce the K10^+^ interscale lineage (orange), and fast-cycling epidermal SCs produce the K36^+^ scale lineage (blue). (**B**) Immunostaining of fibulin-5 (green) in sections of mouse tail skin from 2-month-old versus 30-month-old C57BL/6J mice. The white dashed line represents the epidermal-dermal boundary and hair follicles. Scale bars: 50 µm. (**C**) Immunostaining of fibulin-5 (green) in sections of mouse tail skin from 3-month-old *Fbln5* WT versus KO mice. The white dashed line represents the epidermal-dermal boundary and hair follicles. Scale bars: 50 µm. (**D**) Images of 12-month-old *Fbln5* WT and KO mice. (**E**) The body weight of 12-month-old *Fbln5* WT and KO mice. The data are presented as the mean ± SD. Each dot represents one mouse. The data were compared between groups using a two-tailed *t*-test: *, *P* < 0.05. (**F**–**I**) Hematoxylin and eosin staining of skin sagittal sections of 3- and 12-month-old *Fbln5* WT versus KO mice and quantification (G, I). Scale bars: 150 µm. Epidermal thickness was measured in interscale and scale regions. The data are presented as the mean ± SD. Each dot represents one mouse. The data were compared between groups using a two-tailed *t*-test: **, *P* < 0.005; ns, not significant. (**J**–**M**) Whole-mount staining of BrdU (green, a proliferation marker) and the Hoechst (blue) in 3- and 12-month-old *Fbln5* WT versus KO mice and its quantification (L, M). Scale bars: 200 µm. The data are presented as the mean ± SD. Each dot represents one mouse. The data were compared between groups using a two-tailed *t*-test: **, *P* < 0.005; ns, not significant. (**N-S**) Whole-mount staining of K10 (green, interscale lineage), K36 (red, scale lineage), and the Hoechst (blue) in 2-month-old versus 30-month-old C57BL/6J mice and 3- and 12-month-old *Fbln5* WT versus KO mice and their quantification (O, Q, S). Scale bars: 200 µm. The data are presented as the mean ± SD. Each dot represents one mouse. The data were compared between groups using a two-tailed *t*-test: *, *P* < 0.05; **, *P* < 0.005; ns, not significant.

**FIGURE 2.**
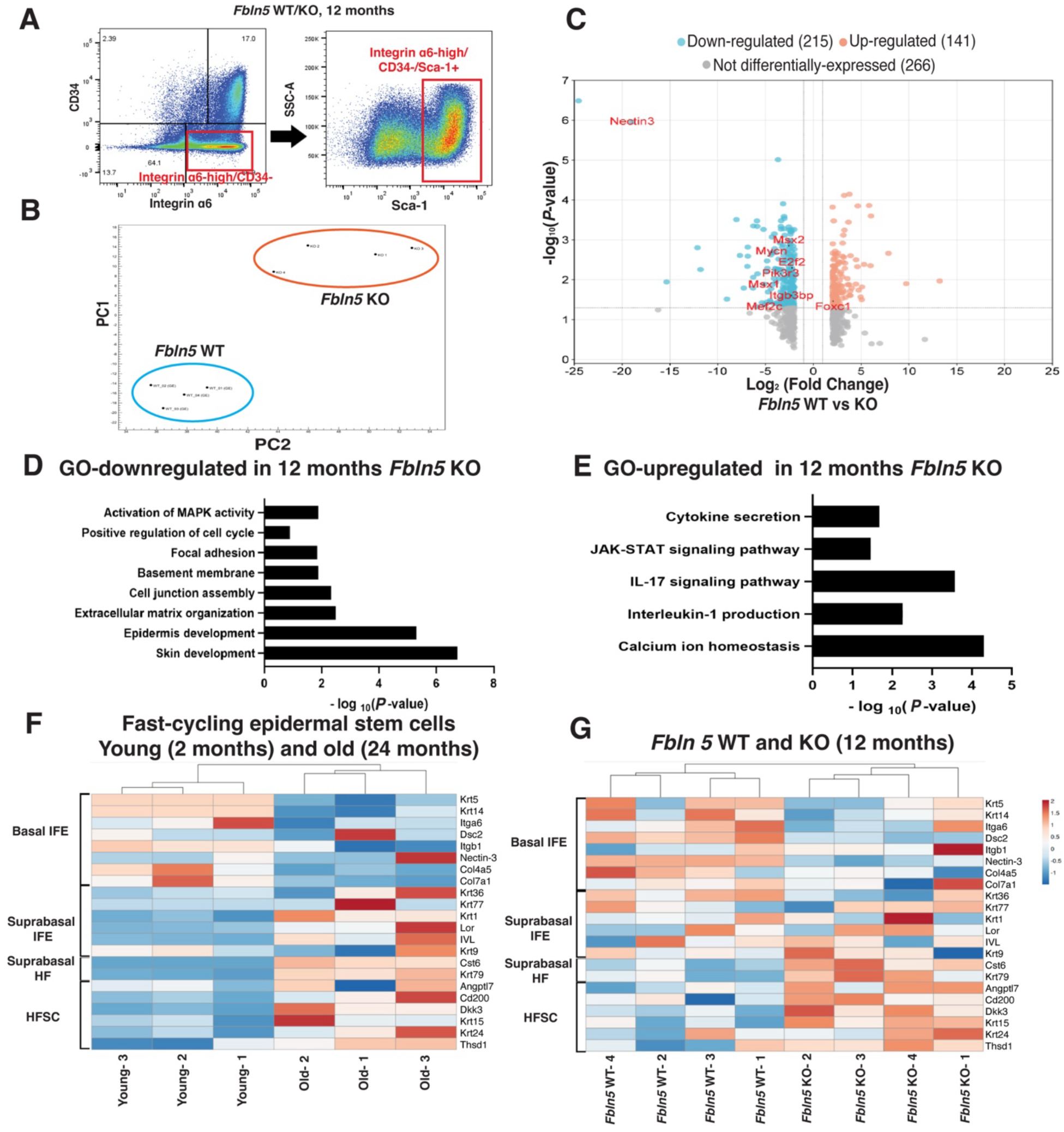
Transcriptome analysis of epidermal stem cells in *Fbln5*-deficient mice. (**A**) Representative FACS plots. Integrin α6^high^/CD34^−^/Sca1^+^ cells were isolated as epidermal stem cells. (**B**) Principal component analysis of twofold DEGs in 12-month-old *Fbln5* WT and KO mice. Each dot represents one mouse (*n* = 4/group). (**C**) Volcano plots of DEGs in 12-month-old *Fbln5* WT and KO mice. (**D, E**) Gene ontology analysis of twofold downregulated (D) and upregulated (E) DEGs in *Fbln5* KO versus WT mice. (**F, G**) The heatmap shows basal and suprabasal signature genes of the interfollicular epidermis and hair follicle lineages based on the RNA-seq data of young (2-month-old) and old (24-month-old) fast-cycling epidermal stem cells and 12-month-old *Fbln5* WT and KO epidermal stem cells.

The length of the Nectin-3^+^ area (Fig. 3T, V; Fig. S3E) and the overlap ratio between Nectin-3 and K36 (Fig. S3F) were measured using the ImageJ software. The YAP, Ki-67, SLC1A3, and ASS1 signals were calculated using the maximum intensities of the nucleus regions and regions adjacent to the nucleus with the ImageJ software (Fig. 4C, E, H, J, L, M, N). More than 100 cells were measured for each experiment.

**FIGURE 3.**
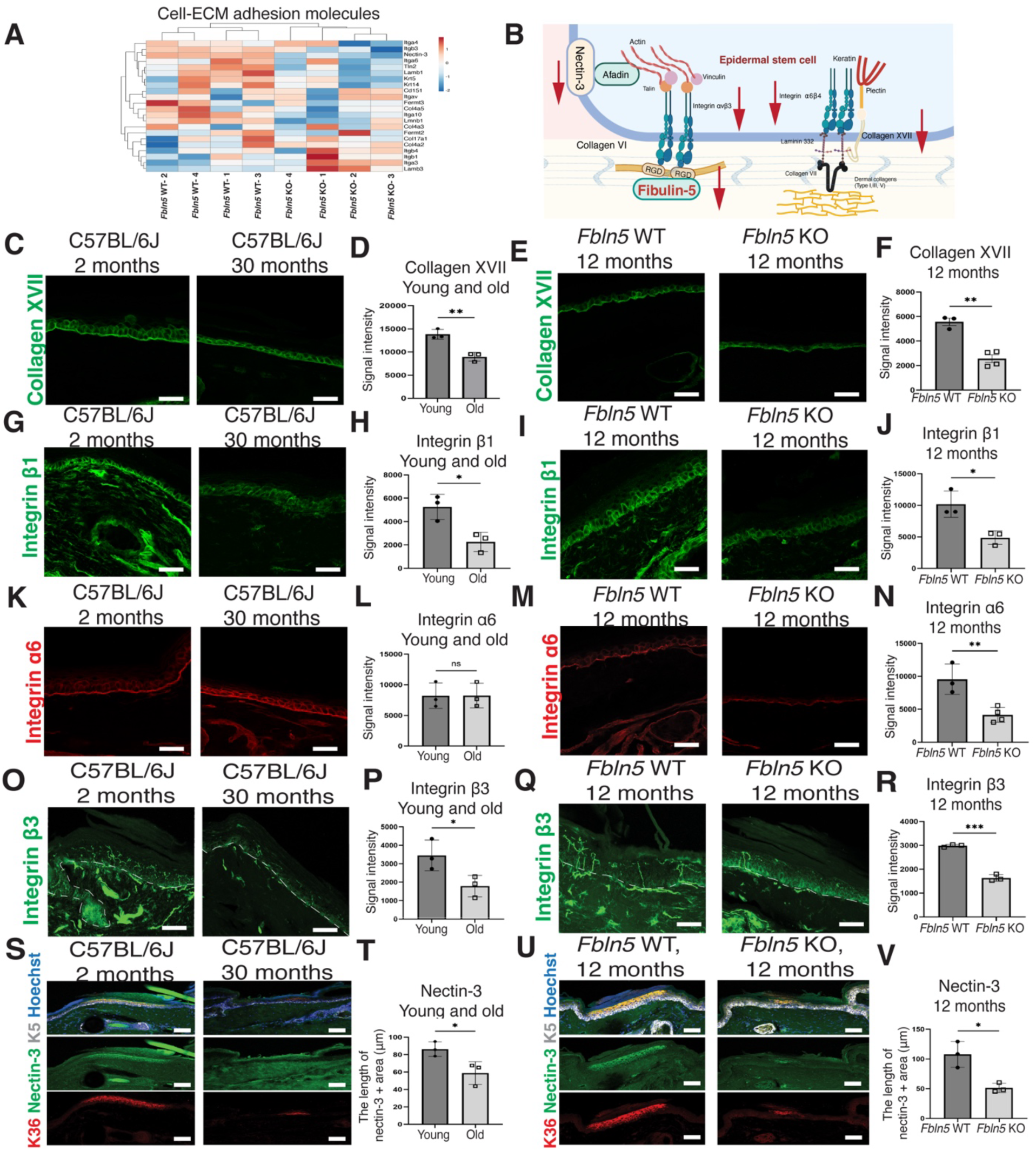
Changes in integrin-extracellular matrix expression due to the loss of fibulin-5. (**A**) The heatmap shows changes in integrins and the ECM proteins in 12-month-old *Fbln5* WT and KO epidermal stem cells. Two-fold DEGs are used for analysis. (**B**) Schematic representation of the epidermal-dermal junction and its associated proteins. (**C**–**V**) Immunostaining of the indicated proteins: collagen type XVII (C, E; green), integrin β1 (G, I; green), integrin α6 (K, M; red), integrin β3 (O, Q; green; scale bar: 20 µm), Nectin-3 (S, U; green), K5 (S, U; grey), and K36 (S, U; red, scale lineage) and their quantification (D, F, H, J, L, N, P, R, T, V). The white dashed line represents the epidermal-dermal boundary and hair follicles. Scale bars: 50 µm. The data are presented as the mean ± SD. Each dot represents one mouse. The data were compared between groups using a two-tailed *t*-test: *, *P* < 0.05; **, *P* < 0.005; ***, *P* < 0.001; ns, not significant. The schematic in panel B was created with BioRender.com.

**FIGURE 4.**
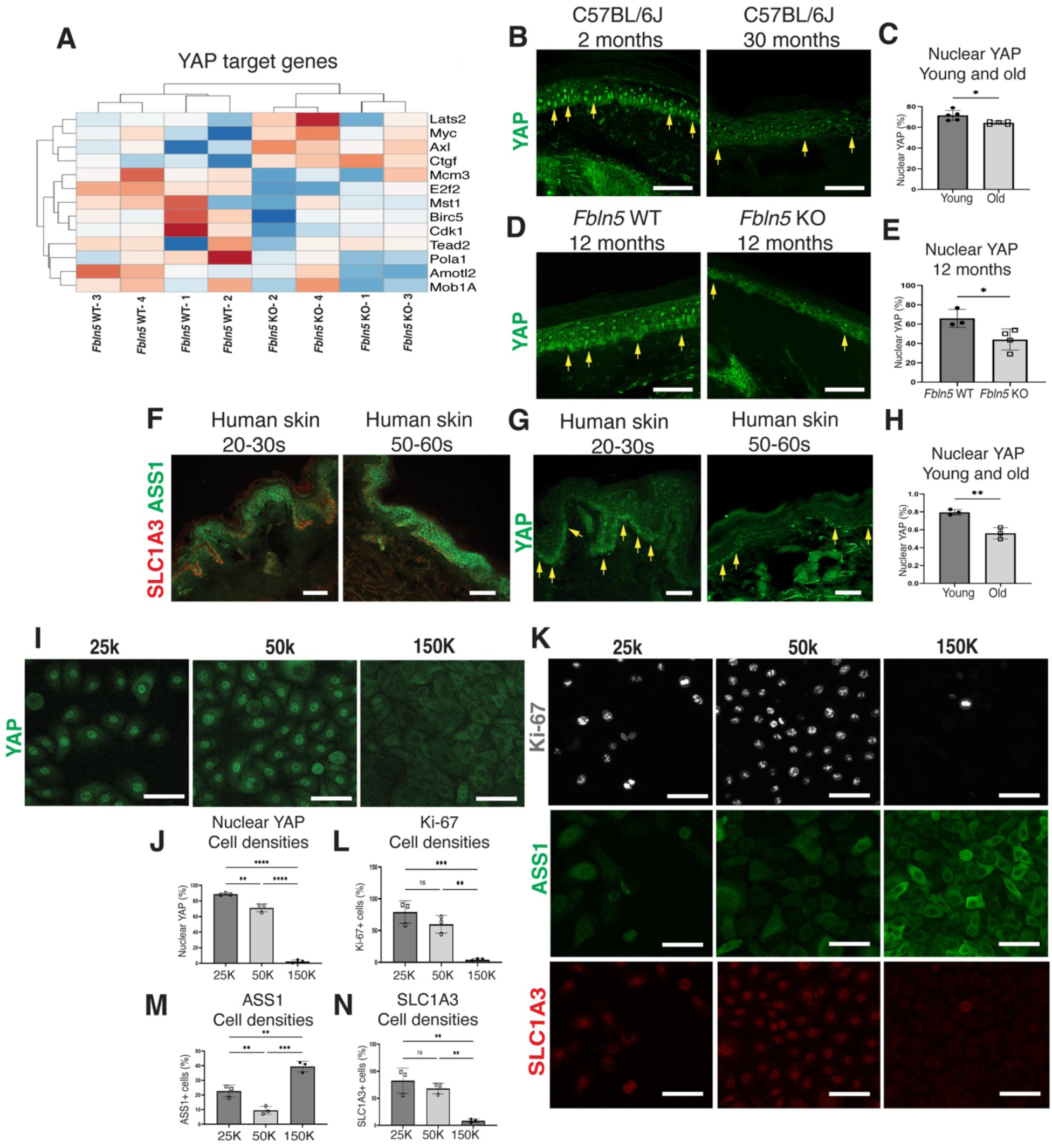
Downregulation of YAP correlates with a decrease in fast-cycling epidermal stem cells. (**A**) The heatmap shows changes in YAP target gene expression in 12-month-old *Fbln5* WT and KO epidermal stem cells; twofold DEGs were used in the analysis. (**B**–**E**) Immunostaining of YAP (green) and the Hoechst (blue) and their quantification (C, E). Scale bars: 50µm. The data are presented as the mean ± SD. Each dot represents one mouse. The data were compared between groups using a two-tailed *t*-test: *, *P* < 0.05. (**F**) Immunostaining of ASS1 (green, slow-cycling epidermal stem cells) and SLC1A3 (red, fast-cycling epidermal stem cells) in sections of young (aged 20–30s years) and old (aged 50–60 years) human skin. Scale bars: 100 µm. The yellow arrows indicate nuclear YAP signals. (**G, H**) Immunostaining of YAP (green) and its quantification (H). Scale bars: 50 µm. The data are presented as the mean ± SD. Each dot represents an individual sample. The data were compared between groups using a two-tailed *t*-test: **, *P* < 0.005. (**I**–**N**) Immunostaining of YAP (I, green), Ki-67 (K, grey), ASS1 (K, green), and SLC1A3 (K, red) in human keratinocytes cultured at different densities and their quantification (J, L, M, N). The cells were seeded at 150,000, 50,000, and 25,000 cells/well in 12-well plates and cultured for 48 h before analysis. Scale bars: 50 µm. The data are presented as the mean ± SD. Each dot represents one experiment. The data were compared between groups using one-way analyses of variance: **, *P* < 0.005; ***, *P* < 0.001; ****, *P* < 0.0001; ns, not significant.

## 3. Results

### 3.1. *Fbln5* KO Mice Showed Early Impairments Resembling Epidermal Stem Cell Aging

To examine age-related changes in *Fbln5* expression in mice, we performed immunostaining on 2-month-old (young) and 30-month-old C57BL/6J mice. In young skin, fibulin-5 was observed around the hair follicles in the dermis and exhibited candelabra-like structures below the interfollicular epidermis (Fig. 1A, B). The absence of dermal signals in *Fbln5* KO mice confirmed the specificity of the fibulin-5 antibody (Fig. 1C). We observed a decrease in fibulin-5 levels in aged skin, particularly in the papillary dermis region below the interfollicular epidermis (Fig. 1B), reflecting the age-dependent decline reported in human skin (Kadoya et al., 2005; Langton et al., 2019).

To assess the impact of fibulin-5 deletion, we analyzed the skin phenotypes of *Fbln5* KO mice. As reported previously, young *Fbln5* KO mice showed saggy skin (Yanagisawa et al., 2002). By 12 months of age, *Fbln5* KO mice exhibited additional gross phenotypes, including a brownish coat, thinner hair, and lower body weight than wild-type (WT) mice in both males and females (Fig. 1D, E). Histologically, epidermal thickness did not differ significantly between mice aged 3 or 6 months (Fig. 1F, G; Fig. S1A, B); however, the epidermis became significantly thinner in the *Fbln5* KO mice compared to the *Fbln5* WT mice by 12 months (Fig. 1H, I). The proliferation of epidermal stem cells also did not differ between 3- and 6-month-old *Fbln5* KO mice but was significantly reduced in 12-month-old mice compared to age-matched control mice (Fig. 1J–M; Fig. S1C–F). These phenotypes indicate age-related skin atrophy, as previously reported (Changarathil et al., 2019).

To further address changes in epidermal stem cells, we examined the dynamics of epidermal stem cells throughout development and aging. The interscale and scale structures in tail skin are formed at the neonatal stage by 2 weeks of age, which coincides with the establishment of epidermal stem cell heterogeneity (Dekoninck et al., 2020; Gomez et al., 2013; Sada et al., 2016). In young adult mice, these structures maintain a constant size, replenished by slow- and fast-cycling epidermal stem cells (Sada et al., 2016). However, the size of the scale regions was significantly reduced in chronologically-aged mice (Fig. 1N, O), consistent with the loss of the fast-cycling epidermal stem cell population (Raja et al., 2022).

In *Fbln5* KO mice, the interscale and scale sizes were unaffected at the neonatal stage (Fig. S1G, H) and in young adults (3 and 6 months; Fig. 1P, Q; Fig. S1I, J). However, by 12 months, while WT mice retained normal interscale/scale size, *Fbln5* KO mice exhibited a significant decrease in scale size (Fig. 1R, S). These results suggest that the loss of *Fbln5* does not significantly affect epidermal stem cell development and homeostasis at a younger age. As the skin ages, the effects of *Fbln5* deficiency on epidermal stem cells become more pronounced, possibly due to chronic inflammation and/or interactions with other ECMs.

### 3.2. *Fbln5* KO Mice Exhibit Molecular Changes Indicative of Epidermal Stem Cell Aging

To explore the molecular changes in epidermal stem cells induced by the loss of *Fbln5*, we performed RNA sequencing (RNA-seq) of fluorescence-activated cell sorted (FACS) epidermal stem cells from *Fbln5* WT and KO mice at 12 months of age (Fig. 2A). Principal component analysis revealed global transcriptome changes between *Fbln5* WT and KO mice (Fig. 2B). We identified differentially expressed genes (DEGs) that were significantly upregulated (orange dots) or downregulated (blue dots) in *Fbln5* KO epidermal stem cells (Fig. 2C). Gene Ontology analysis revealed that the downregulated DEGs were enriched for genes related to the cell cycle, the MAPK signaling pathway, focal adhesion, ECM/basement membrane, and skin development (Fig. 2D). Conversely, the upregulated DEGs were enriched for genes related to pathways associated with cytokine production, the JAK-STAT signaling pathway, and calcium ion homeostasis (Fig. 2E). These gene changes have been reported as signatures of aged epidermal stem cells (Ichijo et al., 2022; Raja et al., 2022).

To assess similarities with chronologically aged skin, we compared the 12-month-old *Fbln5* WT/KO datasets to previously characterized fast-cycling epidermal stem cells in young and aged mice (Fig. 2F; Raja et al., 2022). In 12-month-old *Fbln5* KO mice, the expression of epidermal basal markers tended to decrease, while the expression of epidermal differentiation markers and hair follicle lineage tended to increase (Fig. 2G), reflecting the accelerated epidermal stem cell differentiation and lineage impairment, as previously reported in aging mice (Raja et al., 2022). These results suggest that *Fbln5* KO mice exhibit global molecular changes resembling age-dependent impairment of epidermal stem cell properties.

### 3.3. *Fbln5* Deficiency Compromises ECM Integrity at the Epidermal-Dermal Junction in Aging Skin

Among the ECM proteins at the epidermal-dermal junction (Fujiwara, 2024), the RNA-seq data showed a trend of decreased expression for integrins, collagens, and laminins in the 12-month-old *Fbln5* KO mice (Fig. 3A, B). Immunohistochemically examining these ECM proteins involved in epidermal stem cell regulation revealed that collagen XVII and integrin β1 were downregulated in both *Fbln5* KO and chronologically aged mice (Fig. 3C–J). In contrast, integrin α6 was affected only in *Fbln5* KO mice (Fig. 3K–N).

In the RNA-seq data, we found significant downregulation of integrin β3 and Nectin-3 (Fig. S2A). Fibulin-5 is known to bind integrin αvβ3 via its RGD motif (Kobayashi et al., 2007; Nakamura et al., 2002; Yanagisawa et al., 2009). Nectin-3 is an immunoglobulin-like cell adhesion molecule mainly at the adherens junction and plays a role in epidermal stratification (Mollo et al., 2015; Takahashi et al., 1999; Yoshida et al., 2014). It has been suggested that integrin αvβ3 and Nectin-3 interact through their extracellular regions and play roles in the crosstalk between cell-matrix and cell-cell junctions (Sakamoto et al., 2006; Takai et al., 2008). We examined the spatial localization of integrin β3 and Nectin-3 in young WT skin. Integrin β3 showed a fiber-like pattern localized underneath the epidermis (Fig. 3O), similar to the protein localization of fibulin-5 (Fig. 1B). Nectin-3 was predominantly found in the upper layers at the scale region of the epidermis (Fig. 3S). Integrin β3 and Nectin-3 expression were unaffected in 3-month-old *Fbln5* KO mice (Fig. S2B– E). In aged mice and 12-month-old *Fbln5* KO mice, both integrin β3 and Nectin-3 were downregulated, consistent with a pattern of scale reduction observed with K36 (Fig. 3O–V; Fig. S2F). These results suggest that in the absence of fibulin-5, integrin β3 and/or integrin β1 lose their ligand and anchoring scaffold and are gradually destabilized at the cell-matrix junction together with other extracellular proteins, potentially contributing to the impairment of epidermal stem cell function.

### 3.4. YAP Downregulation is Associated With Reduced Fast-Cycling Stem Cells in *Fbln5* KO Mice and Aging Human Skin

Decreased fibulin-5 and integrins may affect the intracellular signals of epidermal stem cells. In the RNA-seq data, genes associated with mitotic YAP signaling, such as *E2f2* and *Cdk1* (Pattschull et al., 2019), were downregulated in the *Fbln5* KO mice (Fig. 4A). Changes in YAP mRNA levels were confirmed by immunostaining of nuclear YAP, showing that *Fbln5* KO mice, as well as chronologically aged mice, showed reduced YAP activation in epidermal stem cells (Fig. 4B–E).

In humans, fibulin-5 expression declines with age (Kadoya et al., 2005; Langton et al., 2019). Therefore, we investigated whether the epidermal stem cell population changes also occur in humans. In young human skin (aged 20–30 years), the undulating (rete ridge) structure exhibits a heterogeneous distribution of epidermal stem/progenitor markers (Ghuwalewala et al., 2022; Jensen et al., 1999; Legg et al., 2003; Wang et al., 2020; Webb et al., 2004). Among them, we found that ASS1 was enriched at the top of the inter-ridges, whereas SLC1A3, a marker of the fast-cycling population identified in mice and human (Ghuwalewala et al., 2022), was enriched at the base of the rete ridges (Fig. 4F). In aged human skin (age 50–60 years), the undulating structures flattened, SLC1A3 expression decreased, and ASS1 expression generally expanded (Fig. 4F). Furthermore, while nuclear YAP was expressed in the epidermal basal layer in human skin at a younger age, its expression was significantly reduced in human skin at an older age (Fig. 4G, H). These phenotypes are reminiscent of the chronologically aged (30-month-old) or *Fbln5* KO mice, which showed the loss of scale (fast-cycling) regions (Fig. 1N, O, R, S) and inactivation of YAP signaling (Fig. 4B–E).

Given the observed inactivation of YAP with the loss of fibulin-5 or chronological aging, we employed an *in vitro* cell-density model to modulate YAP activity (Panciera et al., 2017). Culturing human primary keratinocytes (epidermal stem/progenitor) at a higher density inactivated YAP and increased differentiation markers while epidermal basal markers remained unchanged (Fig. 4I, J; Fig. S3A, B). Under these conditions, SLC1A3 (a fast-cycling epidermal stem cell or rete ridge marker) and Ki-67 (a proliferation marker) were downregulated, while ASS1 (an inter-ridge marker) was upregulated (Fig. 4K–N; Fig. S3C). These *in vivo* and *in vitro* data indicate that downregulation of YAP correlates with a reduction in fast-cycling stem cell markers in mice and humans; YAP may be a factor that links the reduced integrity of integrin and other ECM proteins due to the loss of fibulin-5 with impaired epidermal stem cell function in aging skin.

## 4. Discussion

The environment surrounding epidermal stem cells changes as the skin ages, but the mechanism linking the extracellular to intracellular changes in stem cells remains unknown. In this study, we found that fibulin-5, a matricellular protein that directly binds to integrins and whose expression decreases with aging, plays a role in maintaining epidermal stem cell heterogeneity (Fig. 5). While the function of fibulin-5 on elastic fiber formation in the skin has been extensively studied, this study demonstrates the novel role of fibulin-5 in providing a protective niche environment for epidermal stem cells in aging skin. Because the loss of fibulin-5 is also seen in chronological aging (Kadoya et al., 2005), photoaging (Langton et al., 2019), and cutis laxa (Gharesouran et al., 2021), fibulin-5-mediated regulation of the epidermal stem cell environment may have clinical relevance in humans.

**FIGURE 5.**
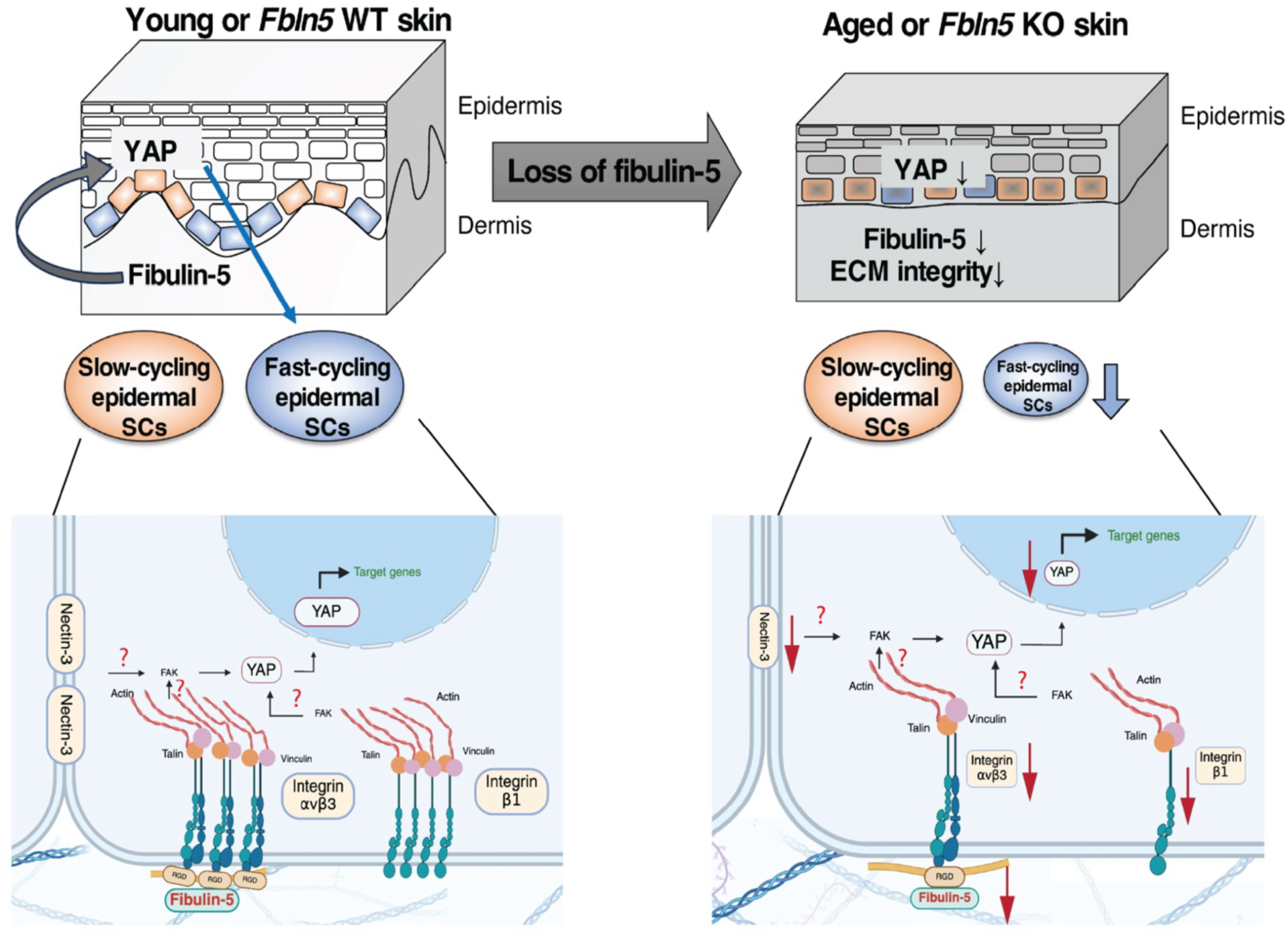
Schematic model of the cellular and molecular changes in aging skin due to fibulin-5 deficiency. In young skin, slow-cycling and fast-cycling epidermal stem cells (SCs) are compartmentalized and produce their respective lineages. An age-dependent decrease in fibulin-5 impairs the expression of integrins and ECM proteins, which alter intracellular signaling, likely through fibulin-5-integrin interactions. In turn, reduced YAP signaling reduces the population of fast-cycling epidermal stem cells in aged human skin and keratinocytes. The schematic was created using BioRender.com.

The loss of fibulin-5 leads to progressive alterations in ECM composition and structure, ultimately disrupting epidermal stem cells’ cellular and molecular properties over time. While compensatory mechanisms may maintain ECM integrity at a young age, these mechanisms remain unclear. As the mice age, there may be combinatorial changes in the ECM, accumulation of DNA damage and mutations, and chronic inflammatory environments in their skin (Jurk et al., 2014; Raja et al., 2022) that exacerbate the changes caused by fibulin-5 deficiency.

While the mechanisms through which fibulin-5 controls epidermal stem cell heterogeneity remain unclear, indirect associations have been suggested. Fibulin-5 binds integrin αvβ3 (Yanagisawa et al., 2009) and integrin β1 (Lomas et al., 2007; Schluterman et al., 2010). Integrin β1 expression was lower in 12-month-old *Fbln5* KO mice (Fig. 3M, N) and human aging skin (Giangreco et al., 2010). Like the *Fbln5* KO and chronologically-aged mice, *Itgb1* KO mice exhibited a loss of scale area (López-Rovira et al., 2005), suggesting a possible integrin β1-fibulin-5 axis in the regulation of epidermal stem cell heterogeneity. Matrix mechanotransduction mediated by integrin-FAK signaling promotes YAP translocation in the skin (Elbediwy et al., 2016) and other tissues (Yamashiro et al., 2020). Cell-cell adhesion proteins also play essential roles in regulating epidermal patterning and YAP activity (Mai et al., 2024). We found that Nectin-3, an adherens junction protein, exhibits spatial heterogeneity, predominantly expressed in scale regions (Fig. 3S). During aging, Nectin-3 expression decreases with the size of the scale region (Fig. 3S, T). Integrin β3, associated with Nectin-3, has also been implicated in regulating the nuclear translocation of YAP *in vitro* (Nardone et al., 2017; Stanton et al., 2019), indicating their possible roles in the mechanotransduction of epidermal stem cells. Further studies are needed to clarify the precise mechanisms of how the cumulative reduction of fibulin-5 affects the extracellular environment and intracellular signaling and subsequently impairs the fast-cycling epidermal stem cells in aging skin.

## Acknowledgments

We thank the Center for Animal Resources and Development at Kumamoto University, the Laboratory of Embryonic and Genetic Engineering at Kyushu University, and the Animal Resource Center at the University of Tsukuba for supporting the animal facilities used in this study. We also thank the International Core-facility of Advanced Life Science at Kumamoto University and the Research Promotion Unit at Kyushu University for their invaluable support. We further thank Dr. Langton (University of Manchester) for her valuable discussion about fibulin-5 in human skin.

Additionally, we thank T. Keida, Y. Kitajima, and K. Ota (Kumamoto University) for their technical assistance with immunostaining and Dr. M. Muratani (University of Tsukuba) for his advice on the RNA-seq analysis. Finally, we thank Dr. A. Yesbolatova (Kyushu University) for the critical reading of this manuscript. This work was supported by the Japan Agency for Medical Research and Development (AMED)—Program for Technological Innovation of Regenerative Medicine (23bm0704067, to AS), AMED-PRIME (21gm6110016, to AS), and Interstellar Initiative Beyond (22jm0610063, to AS)—Grant-in-Aid for Scientific Research (B) from the Japan Society for the Promotion of Science (JSPS) (20H03266 and 24K02035, to AS), Uehara Memorial Foundation (to AS), and The Takeda Foundation (to AS). This work was partly supported by the MEXT Cooperative Research Project Program, Medical Research Center Initiative for High Depth Omics, and CURE (JPMXP1323015486) to the Medical Institute of Bioregulation (MIB), Kyushu University. It was also partly supported by a SPRING grant from the Japan Science and Technology Agency (JST) (JPMJSP2127, to WF).

## Author Contributions

AS conceptualized the project, designed the experiments, and contributed expertise in stem cell and skin biology. WF, MI, ER, AMH, HD, YXN, and YS performed the experiments and analyzed the data. HY provided the *Fbln5* KO mice and fibulin-5 antibodies and contributed expertise on the ECM. YS and KY provided the aged C57BL/6J mice. AS managed and supervised the project. The manuscript was drafted by WF and edited by AS, ER, MI, and HY. Funding was acquired by AS.

## Data Availability

The authors declare that the data supporting the findings of this study are available within the paper and its supplementary information files. The RNA-seq data are deposited in the Gene Expression Omnibus (accession number: GSE295189).

## Declaration of Generative AI and AI-assisted Technologies in the Writing Process

While preparing this manuscript, the authors used ChatGPT to assist with language clarity and readability. Following these tools, the authors thoroughly reviewed and edited the content to ensure its accuracy and take full responsibility for the final version of the published article.

## Supplemental Information

**Supplementary Figure S1.**
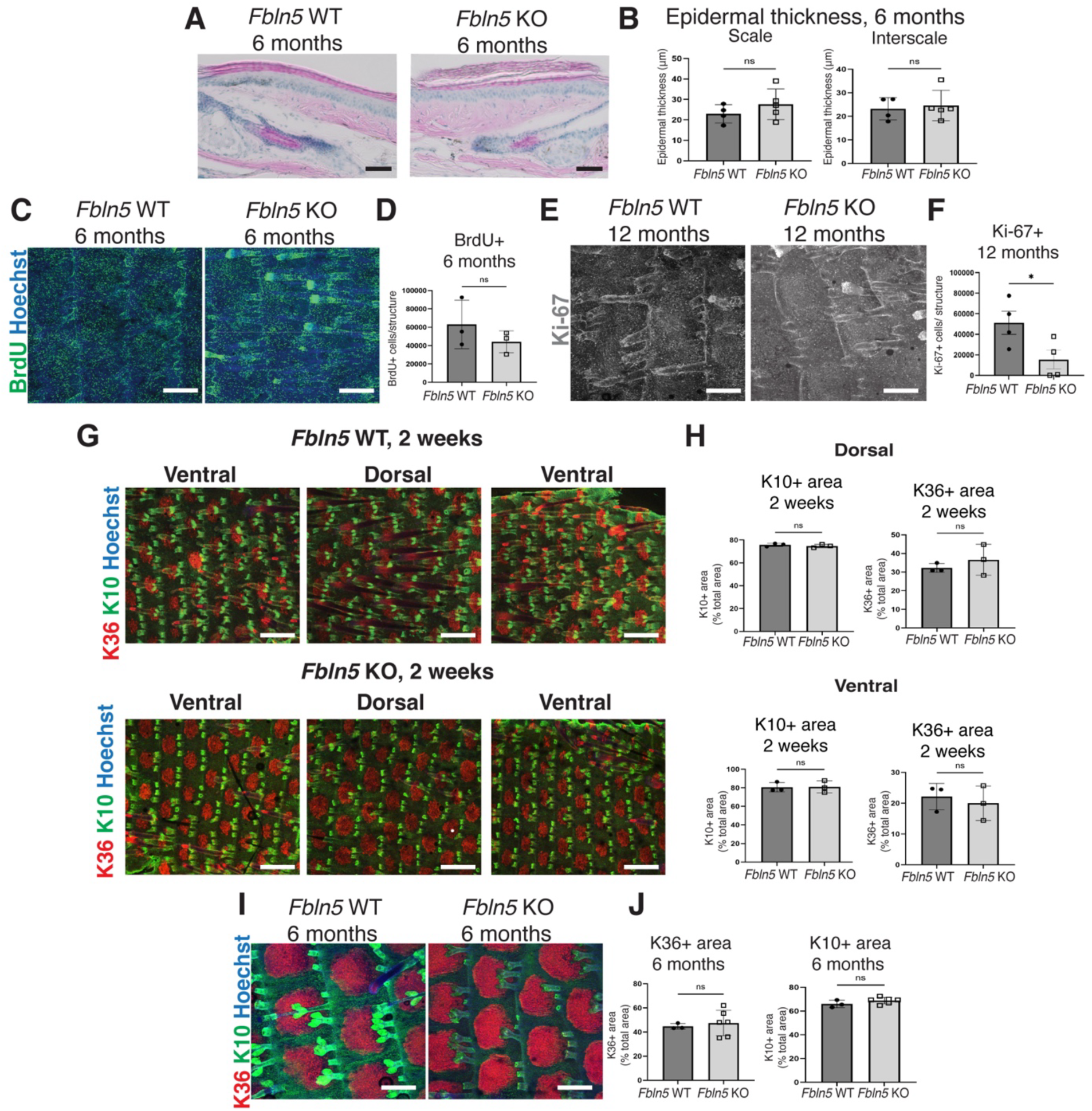
Phenotype analysis of *Fbln5* KO mice at different time points. (**A, B**) Hematoxylin and eosin staining of skin sagittal sections of 6-month-old *Fbln5* WT and KO mice and their quantification (B). Scale bars: 150 µm. Epidermal thickness was measured in interscale and scale regions. The data are presented as the mean ± SD. Each dot represents one mouse. The data were compared between groups using a two-tailed *t*-test: ns, not significant. (**C-F**) Whole-mount staining of BrdU (C, green), the Hoechst (C, blue), Ki-67 (E, grey) from 6- and 12-month-old *Fbln5* WT and KO mice and their quantification (D, F). Scale bars: 200 µm. The data are presented as the mean ± SD. Each dot represents one mouse. The data were compared between groups using a two-tailed *t*-test: *, *P* < 0.05; ns, not significant. (**G, H**) Whole-mount staining of K10 (green, interscale lineage), K36 (red, scale lineage), and the Hoechst (blue) from 2-week-old *Fbln5* WT and KO mice and their quantification (H). Scale bars: 200 µm. The data are presented as the mean ± SD. Each dot represents one mouse. The data were compared between groups using a two-tailed *t*-test: ns, not significant. (**I, J**) Whole-mount staining of K10 (green, interscale lineage) and K36 (red, scale lineage) from 6-month-old *Fbln5* WT and KO mice and their quantification (J). Scale bars: 200 µm. The data are presented as the mean ± SD. Each dot represents one mouse. The data were compared between groups using a two-tailed *t*-test: *, *P* < 0.05; ns, not significant.

**Supplementary Figure S2.**
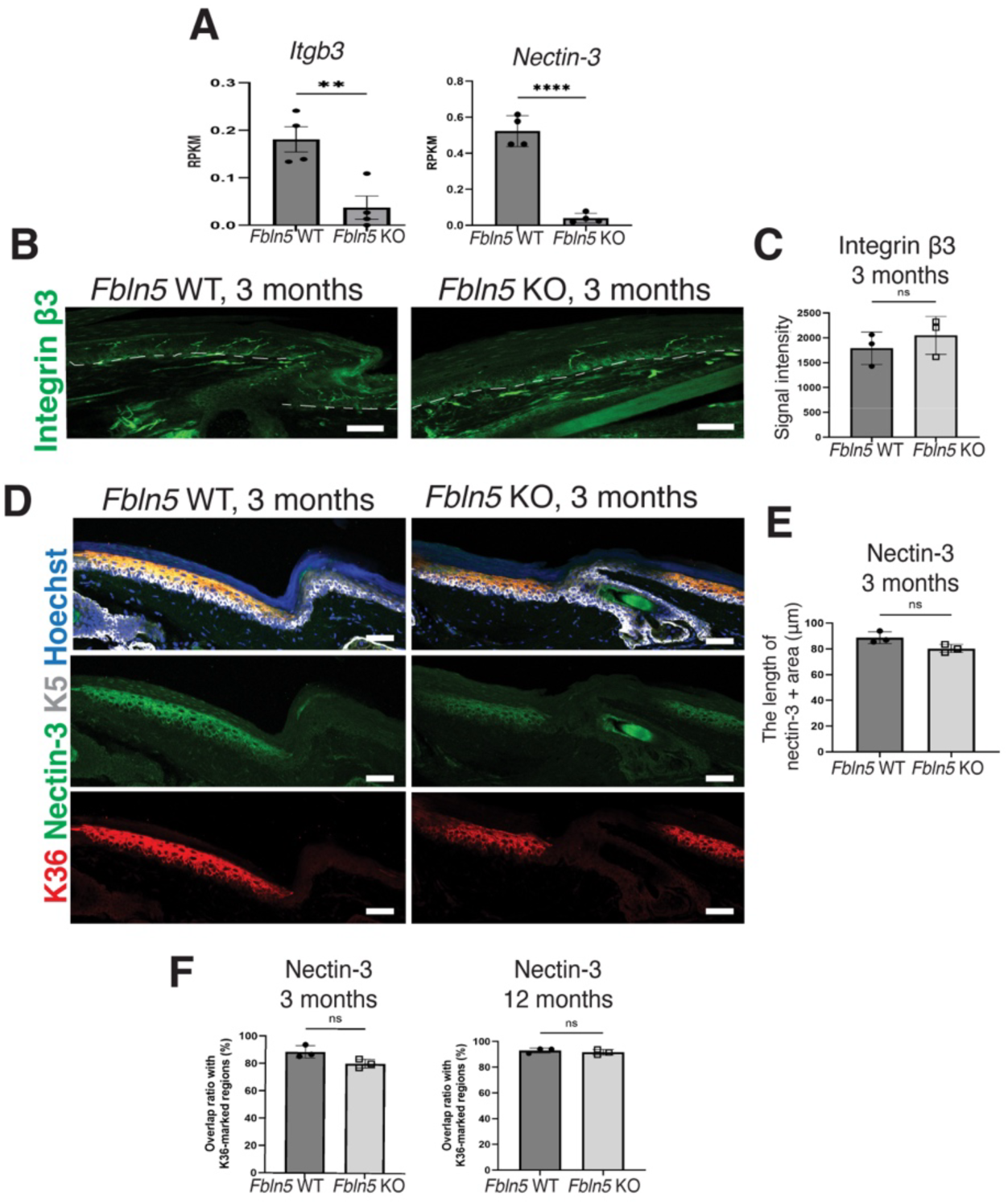
Expression of integrin β3 and Nectin-3 in 3-month-old *Fbln5* KO mice. (**A**) RNA-seq data shows *Nectin3* and *Itgb3* gene expression in 12-month-old *Fbln5* WT and KO epidermal stem cells. The data are presented as the mean ± SD. Each dot represents one mouse. The data were compared between groups using a two-tailed *t*-test: **, *P* < 0.005; ****, *P* < 0.0001. (**B**–**F**) Immunostaining of integrin β3 (B, green, scale bar: 20 µm) and Nectin-3 (D, green, scale bar: 50 µm) and their quantification. The white dashed line represents the epidermal-dermal boundary. The signal intensity of integrin β3 (C), the length of the Nectin-3^+^ area (E), and the overlapping signal of Nectin-3 in K36^+^ regions (F) were quantified. The data are presented as the mean ± SD. Each dot represents one mouse. The data were compared between groups using a two-tailed *t*-test: ns, not significant.

**Supplementary Figure S3.**
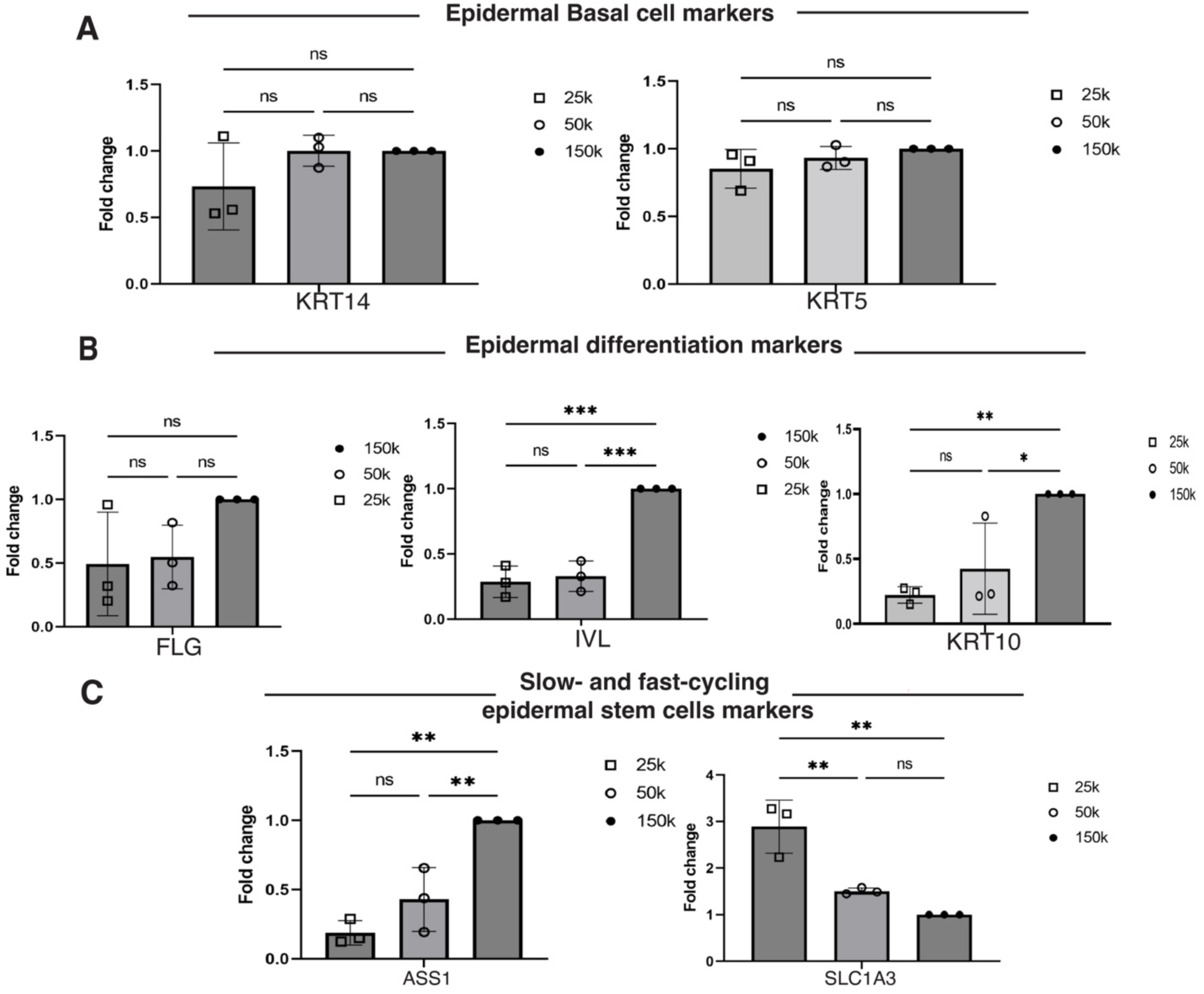
RT-qPCR analysis of human keratinocytes cultured at different densities. (**A**–**C**) RT-qPCR analysis of primary human keratinocytes cultured at different densities normalized to a reference density of 150,000 cells/well. The cells were seeded at 150,000, 50,000, and 25,000 cells/well in 12-well plates and cultured for 48 h before analysis. The data are presented as the mean ± SD. Each dot represents one experiment. The data were compared between groups using one-way analyses of variance: *, *P* < 0.05; **, *P* < 0.005; ***, *P* < 0.001; ns, not significant.

**Supplementary Table 1:**
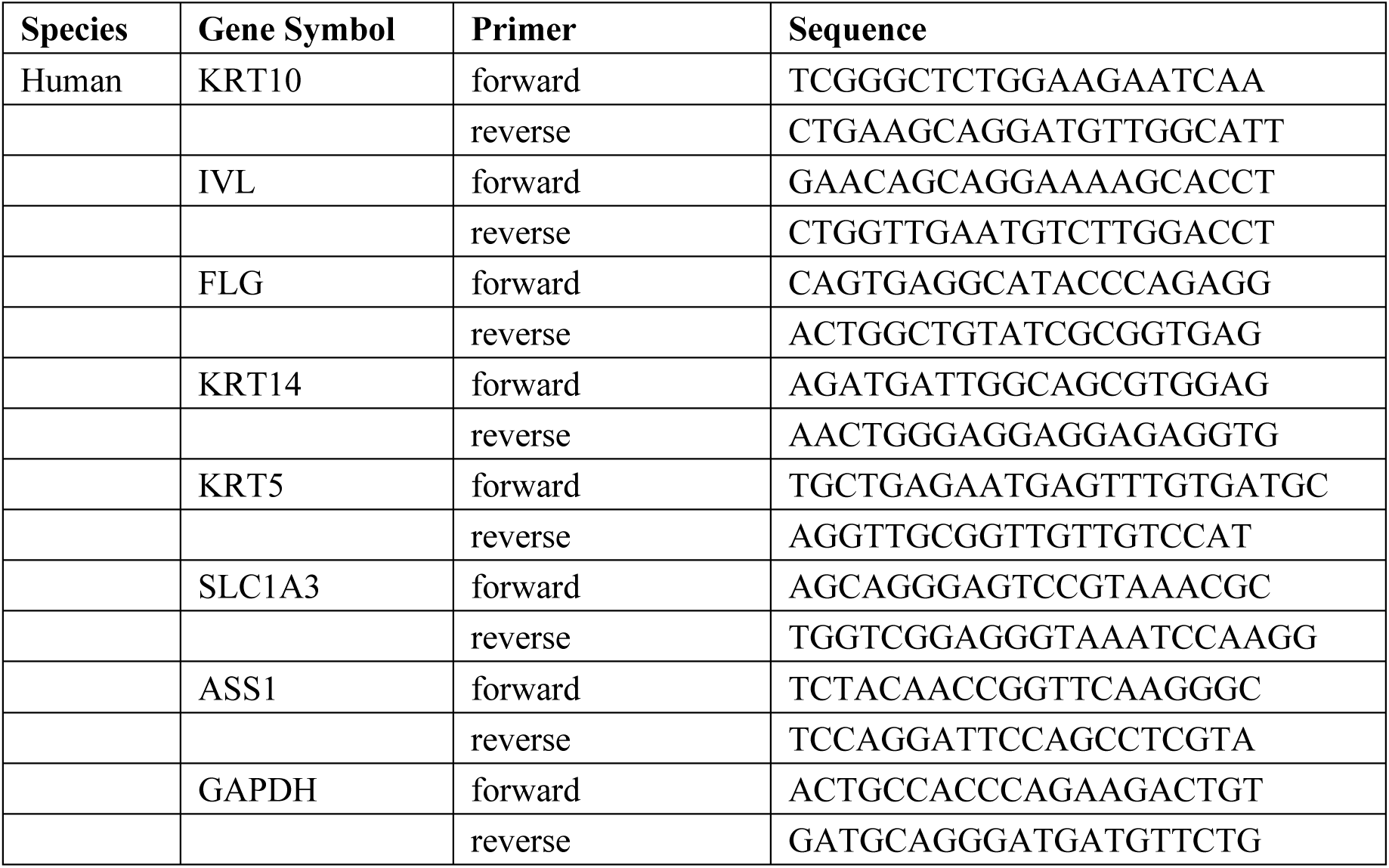
Primers used for qRT-PCR.

## References

AlDahlawi, S., Eslami, A., Häkkinen, L., & Larjava, H. S. (2006). The alphavbeta6 integrin plays a role in compromised epidermal wound healing. Wound repair and regeneration : official publication of the Wound Healing Society [and] the European Tissue Repair Society, 14(3), 289–297. 10.1111/j.1743-6109.2006.00123.x

Aragona, M., Sifrim, A., Malfait, M., Song, Y., Van Herck, J., Dekoninck, S., Gargouri, S., Lapouge, G., Swedlund, B., Dubois, C., Baatsen, P., Vints, K., Han, S., Tissir, F., Voet, T., Simons, B. D., & Blanpain, C. (2020). Mechanisms of stretch-mediated skin expansion at single-cell resolution. Nature, 584(7820), 268–273. 10.1038/s41586-020-2555-7

Budatha, M., Roshanravan, S., Zheng, Q., Weislander, C., Chapman, S. L., Davis, E. C., Starcher, Word, R. A., & Yanagisawa, H. (2011). Extracellular matrix proteases contribute to progression of pelvic organ prolapse in mice and humans. Journal of Clinical Investigation, 121(5), 2048–2059. 10.1172/JCI45636

Changarathil, G., Ramirez, K., Isoda, H., Sada, A., & Yanagisawa, H. (2019). Wild-type and SAMP8 mice show age-dependent changes in distinct stem cell compartments of the interfollicular epidermis. PLOS ONE, 14(5), e0215908. 10.1371/journal.pone.0215908

Chermnykh, E., Kalabusheva, E., & Vorotelyak, E. (2018). Extracellular Matrix as a Regulator of Epidermal Stem Cell Fate. International Journal of Molecular Sciences, 19(4), 1003. 10.3390/ijms19041003

Dekoninck, S., Hannezo, E., Sifrim, A., Miroshnikova, Y. A., Aragona, M., Malfait, M., Gargouri, S., De Neunheuser, C., Dubois, C., Voet, T., Wickström, S. A., Simons, B. D., & Blanpain, C. (2020). Defining the Design Principles of Skin Epidermis Postnatal Growth. Cell, 181(3), 604–620.e22. 10.1016/j.cell.2020.03.015

Elbediwy, A., Vincent-Mistiaen, Z. I., Spencer-Dene, B., Stone, R. K., Boeing, S., Wculek, S. K., Cordero, J., Tan, E. H., Ridgway, R., Brunton, V. G., Sahai, E., Gerhardt, H., Behrens, A., Malanchi, I., Sansom, O. J., & Thompson, B. J. (2016). Integrin signalling regulates YAP/TAZ to control skin homeostasis. Development, dev.133728. 10.1242/dev.133728

Fujiwara H. (2024). Dynamic duo: Cell-extracellular matrix interactions in hair follicle development and regeneration. Developmental biology, 516, 20 – 34. 10.1016/j.ydbio.2024.07.012

Gharesouran, J., Hosseinzadeh, H., Ghafouri-Fard, S., Jabbari Moghadam, Y., Ahmadian Heris, J., Jafari-Rouhi, A. H., Taheri, M., & Rezazadeh, M. (2021). New insight into clinical heterogeneity and inheritance diversity of FBLN5-related cutis laxa. Orphanet Journal of Rare Diseases, 16(1), 51. 10.1186/s13023-021-01696-6

Ghuwalewala, S., Lee, S. A., Jiang, K., Baidya, J., Chovatiya, G., Kaur, P., Shalloway, D., & Tumbar, T. (2022). Binary organization of epidermal basal domains highlights robustness to environmental exposure. The EMBO Journal, 41(18), e110488. 10.15252/embj.2021110488

Giangreco, A., Goldie, S. J., Failla, V., Saintigny, G., & Watt, F. M. (2010). Human Skin Aging Is Associated with Reduced Expression of the Stem Cell Markers β1 Integrin and MCSP. Journal of Investigative Dermatology, 130(2), 604–608. 10.1038/jid.2009.297

Gomez, C., Chua, W., Miremadi, A., Quist, S., Headon, D. J., & Watt, F. M. (2013). The Interfollicular Epidermis of Adult Mouse Tail Comprises Two Distinct Cell Lineages that Are Differentially Regulated by Wnt, Edaradd, and Lrig1. Stem Cell Reports, 1(1), 19 – 27. 10.1016/j.stemcr.2013.04.001

Ichijo, R., Maki, K., Kabata, M., Murata, T., Nagasaka, A., Ishihara, S., Haga, H., Honda, T., Adachi, T., Yamamoto, T., & Toyoshima, F. (2022). Vasculature atrophy causes a stiffened microenvironment that augments epidermal stem cell differentiation in aged skin. Nature Aging, 2(7), 592–600. 10.1038/s43587-022-00244-6

Joost, S., Zeisel, A., Jacob, T., Sun, X., La Manno, G., Lönnerberg, P., Linnarsson, S., & Kasper, M. (2016). Single-Cell Transcriptomics Reveals that Differentiation and Spatial Signatures Shape Epidermal and Hair Follicle Heterogeneity. Cell Systems, 3(3), 221–237.e9. 10.1016/j.cels.2016.08.010

Jurk, D., Wilson, C., Passos, J. F., Oakley, F., Correia-Melo, C., Greaves, L., Saretzki, G., Fox, C., Lawless, C., Anderson, R., Hewitt, G., Pender, S. L., Fullard, N., Nelson, G., Mann, J., Van De Sluis, B., Mann, D. A., & Von Zglinicki, T. (2014). Chronic inflammation induces telomere dysfunction and accelerates ageing in mice. Nature Communications, 5(1), 4172. 10.1038/ncomms5172

Jensen, U. B., Lowell, S., & Watt, F. M. (1999). The spatial relationship between stem cells and their progeny in the basal layer of human epidermis: a new view based on whole-mount labelling and lineage analysis. Development (Cambridge, England), 126(11), 2409 – 2418. 10.1242/dev.126.11.2409

Kadoya, K., Sasaki, T., Kostka, G., Timpl, R., Matsuzaki, K., Kumagai, N., Sakai, L. Y., Nishiyama, T., & Amano, S. (2005). Fibulin-5 deposition in human skin: Decrease with ageing and ultraviolet B exposure and increase in solar elastosis: Age-dependent decrease of fibulin-5 deposition. British Journal of Dermatology, 153(3), 607–612. 10.1111/j.1365-2133.2005.06716.x

Kobayashi, N., Kostka, G., Garbe, J. H. O., Keene, D. R., Bächinger, H. P., Hanisch, F.-G., Markova, D., Tsuda, T., Timpl, R., Chu, M.-L., & Sasaki, T. (2007). A Comparative Analysis of the Fibulin Protein Family. Journal of Biological Chemistry, 282(16), 11805 – 11816. 10.1074/jbc.M611029200

Koester, J., Miroshnikova, Y. A., Ghatak, S., Chacón-Martínez, C. A., Morgner, J., Li, X., Atanassov, I., Altmüller, J., Birk, D. E., Koch, M., Bloch, W., Bartusel, M., Niessen, C. M., Rada-Iglesias, A., & Wickström, S. A. (2021). Niche stiffening compromises hair follicle stem cell potential during ageing by reducing bivalent promoter accessibility. Nature Cell Biology, 23(7), 771–781. 10.1038/s41556-021-00705-x

Koren, E., Feldman, A., Yusupova, M., Kadosh, A., Sedov, E., Ankawa, R., Yosefzon, Y., Nasser, W., Gerstberger, S., Kimel, L. B., Priselac, N., Brown, S., Sharma, S., Gorenc, T., Shalom-Feuerstein, R., Steller, H., Shemesh, T., & Fuchs, Y. (2022). Thy1 marks a distinct population of slow-cycling stem cells in the mouse epidermis. Nature Communications, 13(1), 4628. 10.1038/s41467-022-31629-1

Langton, A. K., Alessi, S., Hann, M., Chien, A. L.-L., Kang, S., Griffiths, C. E. M., & Watson, R. E. B. (2019). Aging in Skin of Color: Disruption to Elastic Fiber Organization Is Detrimental to Skin’ s Biomechanical Function. Journal of Investigative Dermatology, 139(4), 779– 788. 10.1016/j.jid.2018.10.026

Lawlor, K., & Kaur, P. (2015). Dermal Contributions to Human Interfollicular Epidermal Architecture and Self-Renewal. International Journal of Molecular Sciences, 16(12), 28098– 28107. 10.3390/ijms161226078

Legg, J., Jensen, U. B., Broad, S., Leigh, I., & Watt, F. M. (2003). Role of melanoma chondroitin sulphate proteoglycan in patterning stem cells in human interfollicular epidermis. Development, 130(24), 6049–6063. 10.1242/dev.00837

Li, J., Ma, J., Zhang, Q., Gong, H., Gao, D., Wang, Y., Li, B., Li, X., Zheng, H., Wu, Z., Zhu, Y., & Leng, L. (2022). Spatially resolved proteomic map shows that extracellular matrix regulates epidermal growth. Nature Communications, 13(1), 4012. 10.1038/s41467-022-31659-9

Liu, N., Matsumura, H., Kato, T., Ichinose, S., Takada, A., Namiki, T., Asakawa, K., Morinaga, H., Mohri, Y., De Arcangelis, A., Geroges-Labouesse, E., Nanba, D., & Nishimura, E. K. (2019). Stem cell competition orchestrates skin homeostasis and ageing. Nature, 568(7752), 344–350. 10.1038/s41586-019-1085-7

Lomas, A. C., Mellody, K. T., Freeman, L. J., Bax, D. V., Shuttleworth, C. A., & Kielty, C. M. (2007). Fibulin-5 binds human smooth-muscle cells through alpha5beta1 and alpha4beta1 integrins, but does not support receptor activation. The Biochemical Journal, 405(3), 417–428. 10.1042/BJ20070400

López-Rovira, T., Silva-Vargas, V., & Watt, F. M. (2005). Different Consequences of β1 Integrin Deletion in Neonatal and Adult Mouse Epidermis Reveal a Context-Dependent Role of Integrins in Regulating Proliferation, Differentiation, and Intercellular Communication. Journal of Investigative Dermatology, 125(6), 1215 – 1227. 10.1111/j.0022-202X.2005.23956.x

Mai, Y., Kobayashi, Y., Kitahata, H., Seo, T., Nohara, T., Itamoto, S., Mai, S., Kumamoto, J., Nagayama, M., Nishie, W., Ujiie, H., & Natsuga, K. (2024). Patterning in stratified epithelia depends on cell-cell adhesion. Life science alliance, 7(9), e202402893. 10.26508/lsa.202402893

Mascré, G., Dekoninck, S., Drogat, B., Youssef, K. K., Brohée, S., Sotiropoulou, P. A., Simons, B. D., & Blanpain, C. (2012). Distinct contribution of stem and progenitor cells to epidermal maintenance. Nature, 489(7415), 257–262. 10.1038/nature11393

Mollo, M. R., Antonini, D., Mitchell, K., Fortugno, P., Costanzo, A., Dixon, J., Brancati, F., & Missero, C. (2015). P63-dependent and independent mechanisms of nectin-1 and nectin-4 regulation in the epidermis. Experimental Dermatology, 24(2), 114 – 119. 10.1111/exd.12593

Morgner, J., Ghatak, S., Jakobi, T., Dieterich, C., Aumailley, M., & Wickström, S. A. (2015). Integrin-linked kinase regulates the niche of quiescent epidermal stem cells. Nature communications, 6, 8198. 10.1038/ncomms9198

Nakamura, T., Lozano, P. R., Ikeda, Y., Iwanaga, Y., Hinek, A., Minamisawa, S., Cheng, C.-F., Kobuke, K., Dalton, N., Takada, Y., Tashiro, K., Ross Jr, J., Honjo, T., & Chien, K. R. (2002). Fibulin-5/DANCE is essential for elastogenesis in vivo. Nature, 415(6868), 171 – 175. 10.1038/415171a

Nakasaki, M., Hwang, Y., Xie, Y., Kataria, S., Gund, R., Hajam, E. Y., Samuel, R., George, R., Danda, D., M.J., P., Nakamura, T., Shen, Z., Briggs, S., Varghese, S., & Jamora, C. (2015). The matrix protein Fibulin-5 is at the interface of tissue stiffness and inflammation in fibrosis. Nature Communications, 6(1), 8574. 10.1038/ncomms9574

Nardone, G., Oliver-De La Cruz, J., Vrbsky, J., Martini, C., Pribyl, J., Skládal, P., Pešl, M., Caluori, G., Pagliari, S., Martino, F., Maceckova, Z., Hajduch, M., Sanz-Garcia, A., Pugno, N. M., Stokin, G. B., & Forte, G. (2017). YAP regulates cell mechanics by controlling focal adhesion assembly. Nature Communications, 8(1), 15321. 10.1038/ncomms15321

Okuyama, T., Shirakawa, J., Yanagisawa, H., Kyohara, M., Yamazaki, S., Tajima, K., Togashi, Y., & Terauchi, Y. (2017). Identification of the matricellular protein Fibulin-5 as a target molecule of glucokinase-mediated calcineurin/NFAT signaling in pancreatic islets. Scientific reports, 7(1), 2364. 10.1038/s41598-017-02535-0

Panciera, T., Azzolin, L., Cordenonsi, M., & Piccolo, S. (2017). Mechanobiology of YAP and TAZ in physiology and disease. Nature Reviews Molecular Cell Biology, 18(12), 758 – 770. 10.1038/nrm.2017.87

Pattschull, G., Walz, S., Gründl, M., Schwab, M., Rühl, E., Baluapuri, A., Cindric-Vranesic, A., Kneitz, S., Wolf, E., Ade, C. P., Rosenwald, A., Von Eyss, B., & Gaubatz, S. (2019). The Myb-MuvB Complex Is Required for YAP-Dependent Transcription of Mitotic Genes. Cell Reports, 27(12), 3533–3546.e7. 10.1016/j.celrep.2019.05.071

Raja, E., Changarathil, G., Oinam, L., Tsunezumi, J., Ngo, Y. X., Ishii, R., Sasaki, T., Imanaka-Yoshida, K., Yanagisawa, H., & Sada, A. (2022). The extracellular matrix fibulin 7 maintains epidermal stem cell heterogeneity during skin aging. EMBO Reports, 23(12), e55478. 10.15252/embr.202255478

Raja, E., Clarin, M. T. R. D. C., & Yanagisawa, H. (2023). Matricellular Proteins in the Homeostasis, Regeneration, and Aging of Skin. International Journal of Molecular Sciences, 24(18), 14274. 10.3390/ijms241814274

Roca-Cusachs, P., Gauthier, N. C., Del Rio, A., & Sheetz, M. P. (2009). Clustering of α5 β1integrins determines adhesion strength whereas αv β3 and talin enable mechanotransduction. Proceedings of the National Academy of Sciences, 106(38), 16245 – 16250. 10.1073/pnas.0902818106

Rognoni, E., & Watt, F. M. (2018). Skin Cell Heterogeneity in Development, Wound Healing, and Cancer. Trends in Cell Biology, 28(9), 709–722. 10.1016/j.tcb.2018.05.002

Sada, A., Jacob, F., Leung, E., Wang, S., White, B. S., Shalloway, D., & Tumbar, T. (2016). Defining the cellular lineage hierarchy in the interfollicular epidermis of adult skin. Nature Cell Biology, 18(6), 619–631. 10.1038/ncb3359

Sakamoto, Y., Ogita, H., Hirota, T., Kawakatsu, T., Fukuyama, T., Yasumi, M., Kanzaki, N., Ozaki, M., & Takai, Y. (2006). Interaction of Integrin αvβ3 with Nectin. Journal of Biological Chemistry, 281(28), 19631–19644. 10.1074/jbc.M600301200

Stanton, A. E., Tong, X., & Yang, F. (2019). Extracellular matrix type modulates mechanotransduction of stem cells. Acta Biomaterialia, 96, 310 – 320. 10.1016/j.actbio.2019.06.048

Schluterman, M. K., Chapman, S. L., Korpanty, G., Ozumi, K., Fukai, T., Yanagisawa, H., & Brekken, R. A. (2010). Loss of fibulin-5 binding to beta1 integrins inhibits tumor growth by increasing the level of ROS. Disease models & mechanisms, 3(5-6), 333 – 342. 10.1242/dmm.003707

Takahashi, K., Nakanishi, H., Miyahara, M., Mandai, K., Satoh, K., Satoh, A., Nishioka, H., Aoki, J., Nomoto, A., Mizoguchi, A., & Takai, Y. (1999). Nectin/PRR: An Immunoglobulin-like Cell Adhesion Molecule Recruited to Cadherin-based Adherens Junctions through Interaction with Afadin, a PDZ Domain–containing Protein. The Journal of Cell Biology, 145(3), 539–549. 10.1083/jcb.145.3.539

Takai, Y., Miyoshi, J., Ikeda, W., & Ogita, H. (2008). Nectins and nectin-like molecules: Roles in contact inhibition of cell movement and proliferation. Nature Reviews Molecular Cell Biology, 9(8), 603–615. 10.1038/nrm2457

Tang, D., Chen, M., Huang, X., Zhang, G., Zeng, L., Zhang, G., Wu, S., & Wang, Y. (2023). SRplot: A free online platform for data visualization and graphing. PLOS ONE, 18(11), e0294236. 10.1371/journal.pone.0294236

Wang, S., Drummond, M. L., Guerrero-Juarez, C. F., Tarapore, E., MacLean, A. L., Stabell, A. R., Wu, S. C., Gutierrez, G., That, B. T., Benavente, C. A., Nie, Q., & Atwood, S. X. (2020). Single cell transcriptomics of human epidermis identifies basal stem cell transition states. Nature Communications, 11(1), 4239. 10.1038/s41467-020-18075-7

Wang, Y., Kitahata, H., Kosumi, H., Watanabe, M., Fujimura, Y., Takashima, S., Osada, S.-I., Hirose, T., Nishie, W., Nagayama, M., Shimizu, H., & Natsuga, K. (2022). Collagen XVII deficiency alters epidermal patterning. Laboratory Investigation, 102(6), 581 – 588. 10.1038/s41374-022-00738-2

Watt, F. M. (2002). NEW EMBO MEMBER’ S REVIEW: Role of integrins in regulating epidermal adhesion, growth and differentiation. The EMBO Journal, 21(15), 3919 – 3926. 10.1093/emboj/cdf399

Webb, A., Li, A., & Kaur, P. (2004). Location and phenotype of human adult keratinocyte stem cells of the skin. Differentiation, 72(8), 387 – 395. 10.1111/j.1432-0436.2004.07208005.x

Xie, Y., Chen, D., Jiang, K., Song, L., Qian, N., Du, Y., Yang, Y., Wang, F., & Chen, T. (2022). Hair shaft miniaturization causes stem cell depletion through mechanosensory signals mediated by a Piezo1-calcium-TNF-α axis. Cell Stem Cell, 29(1), 70–85.e6. 10.1016/j.stem.2021.09.009

Xuan Ngo, Y., Haga, K., Suzuki, A., Kato, H., Yanagisawa, H., Izumi, K., & Sada, A. (2021). Isolation and Culture of Primary Oral Keratinocytes from the Adult Mouse Palate. Journal of visualized experiments : JoVE, (175), 10.3791/62820. https://doi.org/10.3791/62820

Yamashiro, Y., Thang, B. Q., Ramirez, K., Shin, S. J., Kohata, T., Ohata, S., Nguyen, T. A. V., Ohtsuki, S., Nagayama, K., & Yanagisawa, H. (2020). Matrix mechanotransduction mediated by thrombospondin-1/integrin/YAP in the vascular remodeling. Proceedings of the National Academy of Sciences, 117(18), 9896–9905. 10.1073/pnas.1919702117

Yanagisawa, H., Davis, E. C., Starcher, B. C., Ouchi, T., Yanagisawa, M., Richardson, J. A., & Olson, E. N. (2002). Fibulin-5 is an elastin-binding protein essential for elastic fibre development in vivo. Nature, 415(6868), 168–171. 10.1038/415168a

Yanagisawa, H., Schluterman, M. K., & Brekken, R. A. (2009). Fibulin-5, an integrin-binding matricellular protein: Its function in development and disease. Journal of Cell Communication and Signaling, 3(3–4), 337–347. 10.1007/s12079-009-0065-3

Yoshida, T., Takai, Y., & Thesleff, I. (2014). Cooperation of Nectin-1 and Nectin-3 Is Required for Maintenance of Epidermal Stratification and Proper Hair Shaft Formation in the Mouse. Developmental Biology Journal, 2014, 1–12. 10.1155/2014/432043

